# Depletion of tropomyosin 1.6 promotes matrix-degrading phenotype in TGF-β1-induced myofibroblast

**DOI:** 10.1101/2023.12.08.570775

**Authors:** Ching Lung Wu, Gang Hui Lee, Bo Yu Chen, Cheng Hsiang Kuo, Ting Ting Chan, Yang Kao Wang, Tzyy Yue Wong, Ming Jer Tang

## Abstract

Fibroblasts can be transformed into myofibroblasts under pro-fibrotic conditions, which is characterized by increased contractility and reduced matrix degradation. The relationship between contractile activity and matrix degradation is not fully understood. We found that TGF-β1-induced myofibroblast activation occurs on a culture dish, favoring stress fiber formation and inhibiting podosome structures due to high matrix stiffness. To mimic physiological conditions, we cultured fibroblasts on collagen gel. Blocking actomyosin signaling significantly reduced TGF-β1-induced myofibroblast activation. Tpm1.6, an actomyosin-associated contractile unit, was specifically upregulated by TGF-β1 on soft collagen substrates. Depletion of Tpm1.6 attenuated TGF-β1-induced increase of α-SMA, N-cadherin, and β1-integrin, indicating its crucial role in early myofibroblast activation during fibrosis progression. Tpm1.6 depletion reduced TGF-β1-induced cell contractility and enhanced collagen degradation. Notably, in Tpm1.6-depleted fibroblasts, TGF-β1 triggered formation of distinct α-SMA dot structures enriched with MMP9, promoting collagen degradation. Our study highlights the pivotal role of Tpm1.6 in TGF-β1-induced myofibroblast activation and collagen degradation. Depletion of Tpm1.6 triggers robust collagen degradation through distinct α-SMA dots, presenting a potential therapeutic approach for chronic kidney disease.

## Introduction

End-stage renal disease is characterized by chronic renal inflammation, progressive tubulointerstitial fibrosis, which contributed by myofibroblast activation with α-SMA-mediated contractile phenotype. Myofibroblasts could be originated from various cell sources, including tissue-resident fibroblasts (Kuppe *et al*, 2021), perivascular mesenchymal cells (Humphreys *et al*, 2010; Kuppe *et al*., 2021), and macrophages (Meng *et al*, 2016) during fibrosis. Our previous studies showed that mechanical properties and components of substrates play crucial roles in myofibroblast activation (Chen *et al*, 2014, 2015). However, most *in vitro* studies on myofibroblast activation were conducted on stiff culture dish. Thus, the involvement of the mechanical properties of the microenvironment in myofibroblast activation from different precursors is still needed to be clarified.

Actomyosin-associated contractile units (CUs) are cellular structures responsible for generating contractile forces. For fibroblasts, function of the contractile unit is to exert mechanical forces on surrounding ECM and neighboring cells, and thus plays a crucial role in the developmental process, wound healing and tissue remodeling (Fokkelman *et al*, 2016; Maravillas-Montero & Santos-Argumedo, 2012; Tamada *et al*, 2007). TGF-β1 induce the contractile phenotype of fibroblasts through Rho-ROCK and myosin light chain kinase (MLCK) (Carthy, 2018; Meyer-ter-Vehn *et al*, 2006). Inhibition of actomyosin activity has been shown to suppress fibrosis in lung, liver, and kidney (Nagatoya *et al*, 2002; Southern *et al*, 2016). However, specific CU involved in myofibroblast activation during fibrosis remains unknown. Tropomyosin (Tpm) serves to stabilize actin filaments for regulation of cell motility, adhesion, and vesicle transport via modulation of actomyosin contractility (Gunning *et al*, 2015; Hillberg *et al*, 2006). The Tpm1.6/1.7 isoforms are associated with stress fibers and specifically expressed in fibroblasts, suggesting a crucial role in contractile phenotype of fibroblasts (Gateva *et al*, 2017; Martin & Gunning, 2008; Prunotto *et al*, 2015).

There are two functionally different phenotypes in fibroblasts, i.e. actomyosin contractility and matrix digestion. Activation of Src in epithelial cells suppressed stress fiber and promoted podosome formation (Huveneers *et al*, 2008). Burgstaller and Gomiona demonstrated that cortactin-Arp2/3 microdomain in the podosome leads to the dispersal of myosin/Tpm and reduction of cell contractility via p190RhoGAP (Burgstaller & Gimona, 2004). Those studies supported the notion of opposite and antagonistic relationship between actomyosin contractility and matrix degradation. Under TGF-β1 stimulation, fibroblasts display contractile phenotype even when cultured on collagen gel. In this study, we wished to identify a pivotal CU which contributes to TGF-β1-induced contractility in fibroblasts, and try to reverse phenotype by CU-depletion. We cultured renal fibroblast on collagen gel to mimic initial stage of myofibroblast activation during progression of renal fibrosis, and tried to find a CU component in determining the switch between the contractile phenotype and collagen digestion phenotype induced by TGF-β1 in myofibroblasts. Our study revealed that Tpm1.6 is required for TGF-β1-induced myofibroblast activation, and depletion of Tpm1.6 promotes phenotypic switch from contractility into matrix-degradation upon TGF-β1. Finally, this strong collagen degradation is based on a newly-revealed structure, α-SMA aggregates, which containing MMP.

## Methods

### Cells and reagents

NRK-49F cells (renal fibroblast), primary renal fibroblasts at passage number 6 or below, CCL-226 cells (pericyte), NRK-52E and LLC-PK1 cells (renal epithelium) used for the studies. Cells were tested negative for Mycoplasma assay using TOOLS Mycoplasma Detection Kit (BioTools). Cells were cultured on culture dish, polyacrylamide gel, collagen gel (CG) or gel-coated dish (Co). TGF-β1 (10 ng/ml; PeproTech) is a typical treatment for myofibroblast activation after cell attachment. Primary renal fibroblasts were expanded from the kidney of C57BL/6 mice as described by previous study (Grimwood & Masterson, 2009). Briefly, sections of renal cortex were adhered on gelatin-coated dish and cultured in 1xDMEM/20% FBS condition for 3 days. Fibroblasts proliferated from the explants were cultured and underwent the following experiments under the maximal of six passages. Primary fibroblasts were confirmed by α-SMA and collagen-1a1 staining on culture dish. Detail information of inhibitor treatment and gene silencing are mentioned in the Appendix.

### Western blotting

Protein samples derived from cells were separated by SDS-PAGE and transferred onto PVDF membranes, followed by HRP-conjugated secondary antibodies (CROYEZ) were further used for the recognition of primary antibodies, and visualized by the Pierce ECL Western blot substrate (Thermo Fisher). The intensity of target proteins was analyzed by ImageJ and further normalized with internal control (GAPDH or β-actin for cell lysates; Ponceau S for culture media). The antibodies information is described in the Appendix.

### Immunofluorescence staining

Tissues and cultured cells were fixed by 4% paraformaldehyde, and washed by PBS thrice. Tissue sections and fixed cells were permeabilized by 0.1% Triton X-100, for 15 minutes. Subsequently, washed and blocked by SuperBlock buffer (Thermo Fisher). After staining of primary and secondary anibodies, fluorescence was captured by confocal microscope (FV3000, Olympus Corporation). The antibodies information is described in the Appendix.

### PCR and Real-time qPCR

Total cell RNAs were extracted by TRIzol (Invitrogen), and complementary DNA was synthesized from 1000 ng of total RNA with oligo-dT/random primers and the reverse transcriptase (Thermo Fisher). PCR studies was performed by the amplification of exons or transcripts of Tpm. The cyclic number in traditional PCR is twenty-five. Taqman probes, SYBR Green and StepOne plus real-time PCR machine (Applied Biosystems) were used in real-time PCR studies. Ct values were normalized to β-actin, and relative expression was calculated from 2ΔΔCt calculation. Tpm1.6 and 1.7 were determined independently by using Taqman probe conjugated with 6-FAM-BHQ-1 or TAMRA-BHQ-2. The positive signals of 6-FAM and TAMRA were verified by over-amplification using amplified PCR product. The detailed sequences of primers are mentioned in the Appendix and Appendix Table S1.

### Preparation of polyacrylamide gel

Preparation of polyacrylamide gel was according to Wei et al. (Yeh *et al*, 2010). The of polyacrylamide gels with different stiffness (0.2, 2, 20 kPa) was modified by different the concentration of bisacrylamide and acrylamide with TEMED and ammonium persulfate (Sigma-Aldrich). Detailed procedure is described in the Appendix.

### Preparation of collagen gel

Rat tail type I collagen stock (Corning), 5.7x DMEM, 2.5% NaHCO_3_, 0.1M HEPES, 0.17M CaCl_2_, 1N NaOH and growth medium were mixed together (final collagen concentration: 1 mg/ml), and the mixture was poured on culture dish and waited for polymerization at least for 30 minutes. Collagen gel solution was used for the rinse of the surface of culture dish in the collagen gel-coated dish which is the control condition of collagen gel.

### Culture media precipitation

Harvested culture media derived from cells cultured on soft collagen gel (NRK-49F cells or NRK-49F cells harboring different RNA interference) for 2 days. Removed cell debris by centrifugation at 250g for 5 minutes. Total protein in culture media were precipitated by cold acetone with two-fold volume and further waited for 1 hour at °20 degree. Collected the protein lysates by centrifugation at 13,200g for 10 min and suspend the proteins by sample buffer for SDS-PAGE. Ponceaus S staining was used as the loading control of culture media.

### Micropost array detector (mPAD)

Polydimethylsiloxane (PDMS) micropost arrays were fabricated on cover glass (22mm x 22mm) as previously mentioned (Fu *et al*, 2010). Detailed making procedure of mPAD is described in the Appendix.

### Co-axis system of atomic force microscope and confocal microscope

NRK-49F cells or NRK-49F cells harboring RNA interference were transfected with LifeAct RFP-expressed vector which allowing RFP incorporate with actin filament for the cell position at the live-cell imaging system. LifeAct RFP-expressing cells were cultured on FITC-conjugated collagen gel which can be observed without following staining. The preparation of FITC-conjugated collagen gel is same as collagen gel preparation above, but the collagen stock was conjugated by FITC (Sigma) and further dialysis. After cell culture and treatment under FITC-conjugated collagen gel condition for 4 days, the cell and collagen fibrils were observed and detected by co-axis system of atomic force microscopy and confocal microscopy. After probe calibration and position adjustment, this living *in vitro* system was scanned using optical system that combined laser-scanning confocal microscopy (FV3000, Olympus Corporation) and atomic force microscopy (BioScope Resolve, Bruker Corporation; cantilever: PFQNM-LM-A-CAL), performed as previously mentioned with minor modification (Sandin *et al*, 2020). Briefly, AFM studies were operated in the mode of FASTForce Volume at a scanning rate of 20 Hz. Indentation force was adjusted to maintain 2 nm deformation. Force-distance curves in the indicated area were selected and analyzed with NanoScope Analysis (V1.9, Bruker Corporation) based on Sneddon model. The individual values of cell stiffness and fibril stiffness was displayed on the figures as dots.

### Collagen hybridizing peptide (CHP) recognition

NRK-49F cells or NRK-49F cells harboring RNA interference were cultured on FITC-conjugated collagen gel with or without TGF-β1 induction for 24 h and co-treated with DMSO, blebbistatin (actomyosin blocker, 20 μM; Tocris), and GM6001 (pan-MMP inhibitor, 20 μM; Sigma-Aldrich). The cell-matrix scaffolds were fixed by 4% paraformaldehyde for 1 h and washed by PBS thrice. The 5 μM Cy3-CHP (3Helix) was prepared by heating to 80 °C for 5 min. Heated and cooling Cy3-CHPs were used to apply on fixed cell-matrix scaffolds overnight at 4 °C. After staining, gels were washed with four rounds for 30 min in PBS, followed by the staining of nucleus (Hoechst 33258) and F-actin (Alexa Fluor™ 647 Phalloidin) for cell position. The cell-matrix was observed by confocal microscope (FV3000, Olympus Corporation). The FITC-conjugated collagen fibrils as total fibrils were used for the normalization of collagen deformation. The intensity of CHP-positive fibrils vs. the intensity of FITC-collagen fibrils was analyzed by ImageJ.

### Transmission electron microscope (TEM) inspection

TEM inspection is described in the Appendix.

### Statistics

The GraphPad Prism (version 9.0) software was used for statistical analysis. Unpaired two-tailed Student’s t-tests with Welch’s correction was used to compare the data between two groups; ordinary one-way ANOVA, followed by Tukey’s multiple comparison, was used to compare more than multiple groups; ordinary two-way ANOVA, followed by Tukey’s multiple comparison, was used for cases with multiple variables. The quantitative results were represented as mean with an indication of standard error of the mean (SEM), experimental unit and considered significant (*) with a probability (P) less than 0.05. \**P*<0.05, \*\**P*<0.01, \*\*\**P*<0.001, \*\*\*\**P*<0.0001.

## Results

### Soft matrix shifts contractile phenotype into matrix-degrading phenotype in renal fibroblast

In order to elucidate whether there is correlation between contractile and matrix-degrading phenotype in cultured fibroblasts, we plated renal fibroblasts (NRK-49F) on dish and observed the expression of stress fiber and formation of podosome at different time points. As shown in Figure 1A and 1B, adhered renal fibroblast in culture displayed podosome structure initially at 4 h before the stress fiber was formed. Podosome formation dropped rapidly from 4 h to 12 h, while stress fiber formation was triggered, possibly due to mechanical stimulation from stiff culture dish. At 12 h, the stress fiber and podosome were usually co-expressed in the same cell, but at different localization (Figure 1A). To test whether the matrix stiffness affects the stress fiber and podosome formation, we cultured renal fibroblasts on monomeric collagen-coated polyacrylamide gels with different rigidity (0.2, 2, 20 kPa, and >GPa). The podosome structure was smallest and fewest in cells cultured on a culture dish and showed larger size and greater abundance in cells cultured on softer substrate. Fibroblasts cultured on 0.2kPa displayed highest upregulation of podosome structure (Figure 1C, 1D). The results indicate that there is a mutual exclusiveness between stress fiber and podosome formation and enhanced-matrix stiffness may reduce podosome formation via upregulation of stress fiber and cell contractility in renal fibroblasts.

**Figure 1.**
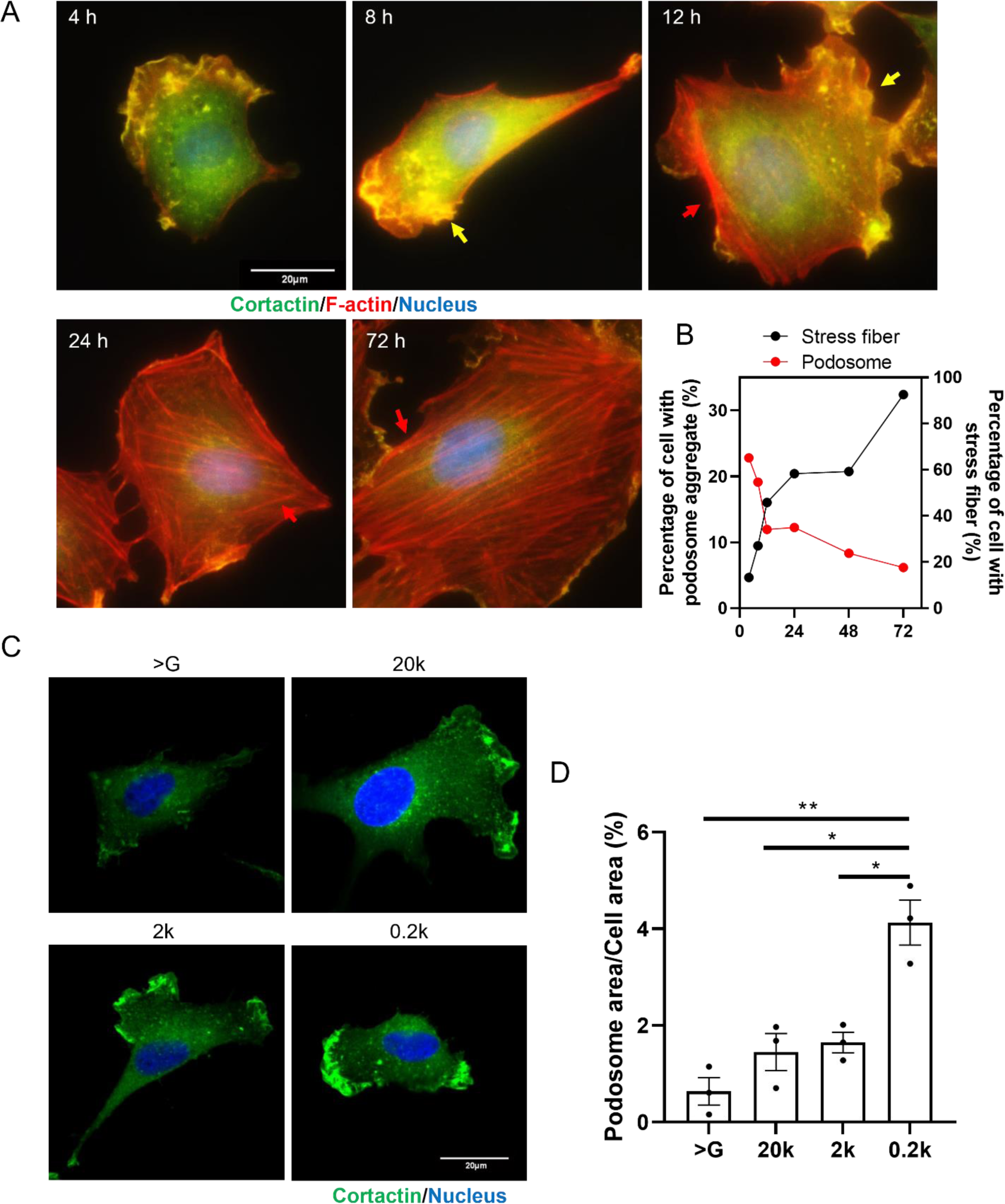
High matrix stiffness of culture dish triggers stress fiber formation and inhibits podosome structures. NRK-49F cells were cultured on regular culture dish for different time (A) or on collagen-coated polyacrylamide gel of different stiffness for 24 h (C) and cultured cells underwent immunofluorescence studies of cortactin (green), phalloidin (red) and nucleus (blue). A. Podosome formation was shown by the localization of podosome structural protein (cortactin) and F-actin staining. Red arrow indicates stress fiber and yellow arrow indicates podosome structure. Scale bar = 20 μm. B. Time course of quantification of podosome- and stress fiber-expressing cells on culture dish for 72 h. C. Fluorescent images show the localization cortactin in fibroblasts cultured on substratum with different stiffness (>G indicates normal culture dish, and 20k, 2k, 0.2k indicate 20kPa, 2kPa, 0.2k Pa. Scale bar = 20 μm. D. Quantification of the area of podosome structure vs. cell area in cells cultured on gel of different matrix stiffness. n=3. Statistical analysis was performed by Welch’s t test.

### TGF-β1-induced myofibroblast activation on collagen gel is observed only in renal fibroblasts

Myofibroblast activation plays a key role in fibrosis progression; however, it is unclear about the origin of cell precursors as well as whether the matrix stiffness affects myofibroblast activation. In renal fibrosis, fibroblasts, macrophages, epithelial cells, and pericytes have been reported to be the precursor of myofibroblasts (Chen *et al*., 2015; Humphreys *et al*., 2010; Wang *et al*, 2016). However, whether these cells can be activated on soft collagen gel (CG), mimicking the physiological interstitial microenvironment, has not been explored before. We examined TGF-β1-induced myofibroblast activation when different types of cells (fibroblasts, pericytes, epithelial cells) were cultured on CG composed of a fibrillar collagen scaffold (<100 Pa). Collagen-gel coated dishes (Co) served as the stiff control, which also contained fibrillar collagen fibrils on the surface. The renal fibroblasts (NRK-49F) showed typical spindle shape morphology on both Co and CG substrates upon TGF-β1 induction, whereas primary renal fibroblasts derived from B6 mouse kidneys cultured on CG displayed less extension, but showed spindle morphology upon TGF-β1 treatment (Appendix Fig S1A, 1B). Both renal fibroblasts cultured on CG showed the phenotype of TGF-β1-stimulated myofibroblast activation, as manifested by elevation of α-SMA, N-cadherin and β1 integrin (Figure 2A-D, 2I). However, TGF-β1 treatment could not induce increase in α-SMA level in pericytes (CCL226) cultured on CG (Appendix Fig S1C-E). Additionally, renal epithelial cells (NRK-52E and LLC-PK1) exhibited limited cell growth and elevated apoptosis on soft CG. TGF-β1 treatment particularly enhanced apoptosis in LLC-PK1 cells cultured on CG (Appendix Fig S1F, 1I). We found that TGF-β1-induced epithelium-to-mesenchymal transition, as reflected by augmentation of α-SMA levels, was significantly ameliorated by CG (Appendix Fig S1G, 1H, 1J, 1K). If fibroblasts were cultured on monomeric collagen-coated polyacrylamide gels with a rigidity of 0.2 kPa, TGF-β1-induced myofibroblast activation was blocked (Appendix Fig S1L, 1M). A matrix which can be remodeled is essential for myofibroblast activation. Among the tested cell types, only fibroblasts retained the ability of TGF-β1-induced myofibroblast activation on CG, suggesting that fibroblasts are the potential precursor of myofibroblasts during onset of fibrosis when matrix stiffness is low.

**Figure 2.**
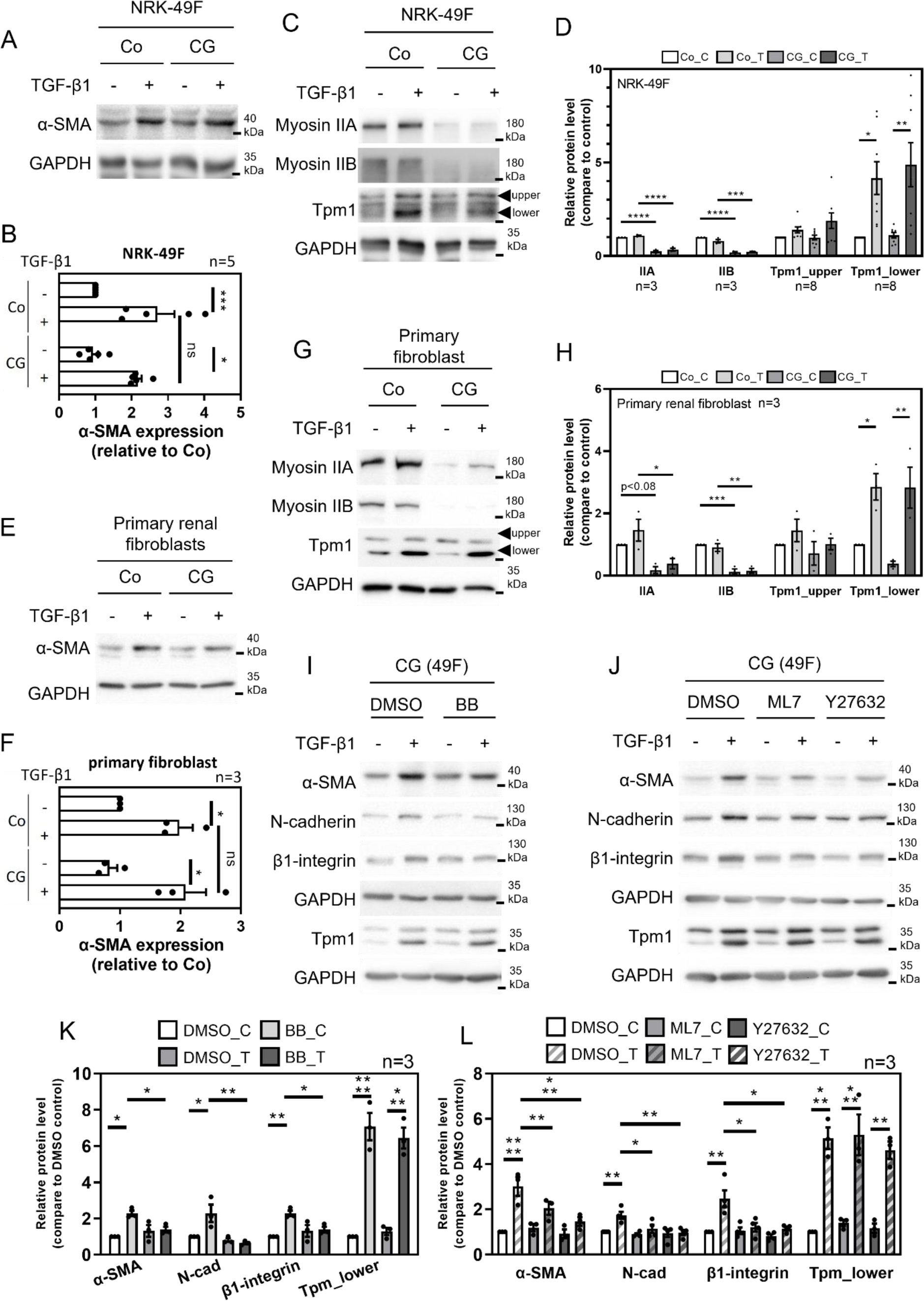
Actomyosin contractility is required for TGF-β1-induced myofibroblast activation under soft collagen gel. A-H. NRK-49F cells and primary renal fibroblast cells were cultured on collagen gel-coated dish (Co) and collagen gel (CG) with and without TGF-β1 (10 ng/ml) for 24 h. The protein levels of myofibroblast marker (α-SMA) and actomyosin-associated contractile units (CUs; myosin IIA, IIB, and Tpm1) in NRK-49F cells (A, C) and primary renal fibroblasts (E, G) were analyzed by Western blotting and quantification results of Western blotting are shown in figure B, D, F, and H. The arrow shows the different isoforms of Tpm1 in NRK-49F cells and primary fibroblasts. Statistical analysis was performed by two-way ANOVA. I-L. Protein levels of α-SMA, N-cadherin, β1-integrin, and Tpm1 in NRK-49F cultured on CG with or without co-treatment of actomyosin blockers (BB: blebbistatin, ML7, Y27632) for 24 h, and the respective quantification results are shown in figure K and L. n=3. Statistical analysis was performed by two-way ANOVA.

### Tropomyosin-1 (Tpm1) expression is elevated in TGF-β1-induced fibroblast-to-myofibroblast activation

Since actomyosin contractibility may be accompanied with myofibroblast activation, we wished to investigate the role of actomyosin contractility in myofibroblast activation when cells were cultured on CG. We examined the levels of myosin IIA/IIB and Tpm1 in NRK-49F cells and primary renal fibroblasts cultured under Co or CG conditions with or without TGF-β1 treatment. Myosin IIA and IIB, but not Tpm1 as recognized by exon 1a-specific antibody (TM311), were downregulated when cells were cultured on CG. At our hand, TM311 recognized Tpm2.1 and different isoforms of Tpm1, as indicated by upper and lower bands, respectively (Figure 2C, 2G). Tpm1 was enhanced by TGF-β1 in NRK-49F cells and primary renal fibroblasts cultured on both Co and CG (Figure 2C, 2D, 2G, 2H). Immunofluorescence staining showed that partial Tpm1 was recruited into stress fiber after TGF-β1 stimulation (Appendix Fig S2A). TGF-β1-induced increase in Tpm1 as well as α-SMA level was ameliorated by inhibitor of TGFβRI (SB431542) and smad3 (SIS3) in NRK-49F cells (Appendix Fig S2B-D). To assess the importance of actomyosin-mediated contractile force in TGF-β1-induced myofibroblast activation, actomyosin activity blockers (blebbistatin, ML7 and Y27632) were applied on NRK-49F cells cultured on Co or CG upon TGF-β1. The results showed that actomyosin inhibition suppressed TGF-β1-induced myofibroblast activation (Figure 2I-L; Appendix Fig S2E, 2F). Interestingly, treatment of actomyosin inhibitors did not alter TGF-β1-induced elevation of Tpm1. Our results indicate that the actomyosin contractility is required for TGF-β1-induced myofibroblast activation.

### TGF-β1 specifically increases the expression of Tpm1 isoform 1.6 in renal fibroblasts

There are various isoforms of Tpm1 due to alternative promoters and splicing sites (Figures 3A). We first analyzed specific exons of Tpm1 exon by real-time PCR (exon 1a, 2a, 2b, 9a, 9d). Exon 1a, 2a, 2b, 9a and 9d of Tpm1 were elevated by TGF-β1 in NRK-49F cells under CG condition, whereas exon 9d of Tpm3 was not altered (Appendix Fig S3A). The protein level of Tpm3 was also not altered by TGF-β1 stimulation (Appendix Fig S3B). To characterize the specific isoform of Tpm1 involved in TGF-β1-induced myofibroblast activation, we analyzed different Tpm1 isoforms by combination of different exon primer sets (e.g. forward primer of exon 1a in combination with reverse primer of exon 9d). The amplicon of paired primer for 1a to 9d and 2b to 9d, but not 1a to 9a, 1b to 9d, or 2a to 9d, was significantly upregulated by TGF-β1 in both NRK-49F cells and primary renal fibroblasts cultured on CG, indicating that the TGF-β1-induced elevated Tpm isoforms are Tpm1.6/1.7 (Figure 3B, 3C, Appendix Fig S3C, 3D). Moreover, we employed two Taqman probes conjugated with different fluorescence reporters to distinguish Tpm1.6 (6FAM-exon 6b targeted sequence-BHQ1) and Tpm1.7 (TAMRA-exon 6a targeted sequence-BHQ2) in total RNA extracts (Figure 3D). The results showed that PCR amplification of 1a to 9d and 2b to 9d released solely the 6FAM into reaction mixtures, which was increased by TGF-β1 treatment (Figure 3E, 3F). The results of exon 1a-initiated Sanger sequencing also demonstrated the presence of only mRNA sequence of Tpm1.6 in NRK-49F cells with or without TGF-β1 treatment (Appendix Fig S3E, 3F). Taken together, Tpm1.6 is the dominant Tpm isoform specifically upregulated by TGF-β1 in renal fibroblasts.

**Figure 3.**
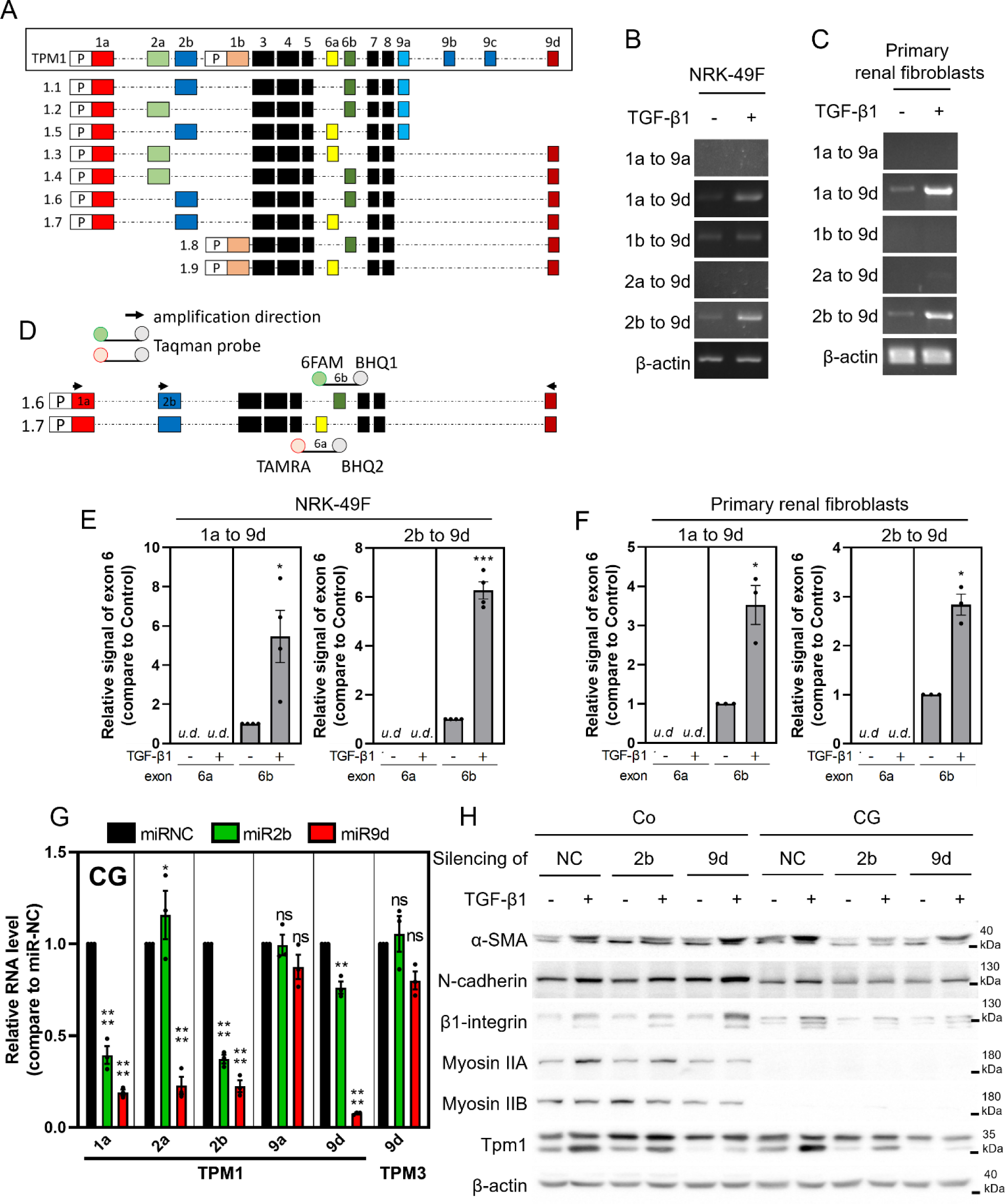
Silencing of the renal fibroblast-dominant isoform, Tpm1.6, suppresses TGF-β1-induced myofibroblast activation. NRK-49F cells and primary renal fibroblasts were cultured on collagen gel (CG) with or without TGF-β1 for 24 h, and total RNA was extracted and analyzed by traditional PCR (B, C) or real-time PCR (E, F). A. Schematic graph displays Tpm1 isoforms derived from alternative promoters and splicing exons. B, C. PCR results represent the levels of different transcripts of Tpm1 in NRK-49F cells (B) and primary renal fibroblasts (C). Isoform1.6/1.7 of Tpm1 (Tpm1.6/1.7) was specifically upregulated by TGF-β1 in renal fibroblasts. n=3. D. Forward/reverse primers and Taqman probes-conjugated with fluorescent reporter were used for quantitative real-time PCR studies in order to detect transcript of specific Tpm1.6 and Tpm1.7 isoforms. The strategy was to recognize Tpm1.6 and 1.7 by 6FAM-BHQ1- and TAMRA-BHQ2-conjugated Taqman probes against exon 6b and 6a, respectively. The polymerase in Tpm1.6/1.7 amplification (1a to 9d or 2b to 9d) passes the exon 6 and releases the reporter fluorescence from Taqman probes by nucleotides cleavage. E, F. Quantitative real-time PCR studies show the two distinct exon 6 signals (6a from TAMRA and 6b from 6FAM) by Tpm1.6/1.7 amplifications (1a to 9d or 2b to 9d) in NRK-49F cells (E; n=4) or primary renal fibroblasts (F; n=3). The level of Tpm1.6, the dominant isoform in renal fibroblasts, is specifically upregulated by TGF-β1. G, H. NRK-49F cells received liposome carried biogenic RNA interference against exon 2b and 9d of Tpm1, and then were cultured on a gel coated-dish (Co) or CG with or without TGF-β1 induction for 24 h. Total RNA was extracted and analyzed by real-time PCR (G), whereas cell lysates were analyzed by Western blotting (H). Quantitative real-time PCR shows the RNA levels of individual Tpm1 exon in NRK-49F cells harboring biogenic RNA interference against exon 2b and 9d of Tpm1. miRNC indicates non-silencing control and miR2b/miR9d indicate specific exon silencing groups. The results indicate that exon 2b-targeting silencing shows more specificity for Tpm1.6. n=3. Western blot represents the protein levels of myofibroblast proteins (α-SMA, N-cadherin, β1-integrin) and contractile units (myosin IIA, IIB, and Tpm1) in NRK-49F cells harboring biogenic RNA interference against exon 2b and 9d of Tpm1. Statistical analysis was performed by Welch’s t test (E, F) or one-way ANOVA (G).

### The silencing of Tpm1.6 suppresses TGF-β1-induced myofibroblast activation and promotes collagen gel degradation

To evaluate the functional role of Tpm1.6 in TGF-β1-induced myofibroblast activation, Tpm1.6 was silenced by liposomes-carried biogenic RNA interference against exon 2b and 9d of Tpm1 in NRK-49F cells. Real-time PCR showed specific inhibition of exon 2b or 9d in cells harboring miRNA mimetic against exon 2b or 9d, respectively (miR2b cells or miR9d cells). Results indicate that exon 2b silencing is more specific for depletion of Tpm1.6, whereas exon 9d silencing may reduce not only Tpm 1.6, but also Tpm 1.3/1.4 which contains exon 2a (Figure 3G). Silencing of exon 2b or 9d did not alter the TGF-β1-induced increase in protein levels of myofibroblast markers in renal fibroblasts cultured on Co, whereas on CG, the depletion significantly reduced the TGF-β1-induced increase in myofibroblast proteins (Figure 3H; Appendix Fig S4A). However, exon 2b silencing exhibited a better effect than exon 9d. Furthermore, the silencing did not alter cell proliferation which was confirmed by Cell Counting Kit-8 assay (Appendix Fig S4B). Taken together, depletion of Tpm1.6 via exon 2b-targeting inhibits TGF-β1-induced myofibroblast activation under CG condition.

To test if the silencing of Tpm1.6 affects cell contractility, collagen contraction assay was employed. Tpm1.6-silenced NRK-49F cells were cultured in CG with or without TGF-β1, and the CGs were released for indicated time to visualize the change of gel area. We found that cells harboring silencing of exon 2b showed decreased amount of gel volume at 4 h. Because the gel contraction was not accompanied with increased gel opaqueness and there was also strong degradation of gel observed at 48 h, such results suggest that Tpm1.6-silenced cells display an active collagen degradation capability upon TGF-β1 induction (Figure 4A, 4B). The traditional gel contraction assay may not be suitable for cell contractility measurement particularly when cultured cells display high matrix-degradation capability.

**Figure 4.**
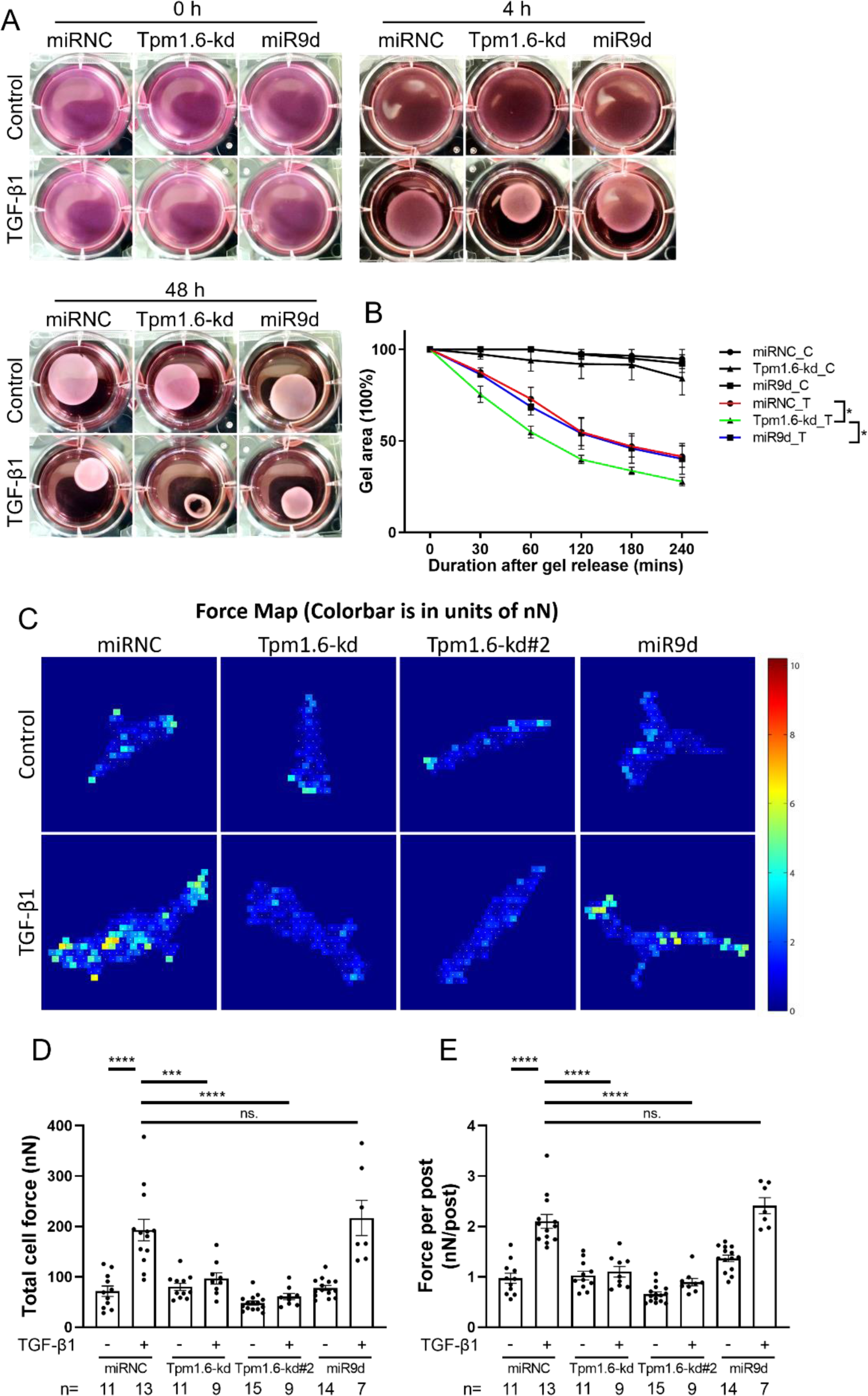
Silencing of Tpm1.6 by exon 2b-targeted RNA interference prevents TGF-β1-induced cell contractility. A, B. NRK-49F cells harboring biogenic RNA interference against Tpm1 were cultured in collagen gel with or without TGF-β1 for 24 h and the collagen gels were released for indicated time to visualize the gel contraction/degradation, and the quantification results are shown in figure B. n=3. Cells harboring silencing of exon 2b shows decreased amount of gel volume without increase opaqueness at 4 h and strong degradation of gel at 48 h, indicating an active process of gel digestion is present. C-E. Cell contractility is measured by micropost array detector. NRK-49F cells harboring biogenic RNA interference against Tpm1 were cultured on PDMS-based microposts with or without TGF-β1 for 24 h. Each post bending in response to applied traction force as calculated by finite element model analysis. The force map represents contractility distribution of non-silencing control and Tpm1.6-silenced cells, and color bar shows the force level (nN). The total cell force and force per post (nN) were quantified and shown in figure D and E. Statistical analysis was performed by RM one-way ANOVA (B) and two-way ANOVA (D, E).

### Tpm1.6 depletion reduces TGF-β1-induced cell contractility

To assess cell contractility, we employed mPAD. Results showed that silenced-control cells exhibited stronger contractile force around cell protrusions in presence of TGF-β1 (71.55 nN to 192.8 nN). Knockdown of Tpm1.6 (Tpm1.6-kd) did not change cell contractility (80.14nN), but markedly attenuated TGF-β1-induced increase in cell contractility (96.54 nN). In contrast, exon 9d-silencing did not prevent TGF-β1-induced increase in cell contractility (Figure 4C, 4D). The suppression of TGF-β1-induced cell contractility in Tpm1.6-kd group was confirmed by another targeting sequence of exon 2b (Tpm1.6-kd#2).

In order to assess the stiffness of single collagen fibril exerted by cell contractility, we employed an advanced technology: confocal microscope co-axis with atomic force microscope (AFM). LifeAct-RFP-transfected cells (red) were cultured on FITC-conjugated CGs (green) for confocal microscopy-based visualization. The structure and rigidity of cells and peri-cellular fibrils were analyzed by AFM. TGF-β1 treatment promoted more fibrils recruitment toward to cell, but did not change the diameter of collagen fibrils (Figure 5A). The stiffness of silenced control cells and fibrils was 8.7±2.3 KPa and 8.3±1.3 KPa, respectively. TGF-β1 markedly enhanced cell stiffness to 32.4±6.4 KPa, as well as the stiffness of fibril exerted by cells (40.7±3.3 KPa; Figure 5B, 5C). Silenced cells did not alter stiffness of cell and collagen fibril. However, Tpm1.6-kd significantly reduced TGF-β1-induced increase in stiffness of cell and collagen fibril. Exon 9d-silencing did not affect TGF-β1-indcued increase in cell rigidity, but slightly attenuated the stiffness of peri-cellular fibrils. Taken the results of mPAD and AFM, Tpm1.6-depletion markedly reduced TGF-β1-induced cell contractility.

**Figure 5.**
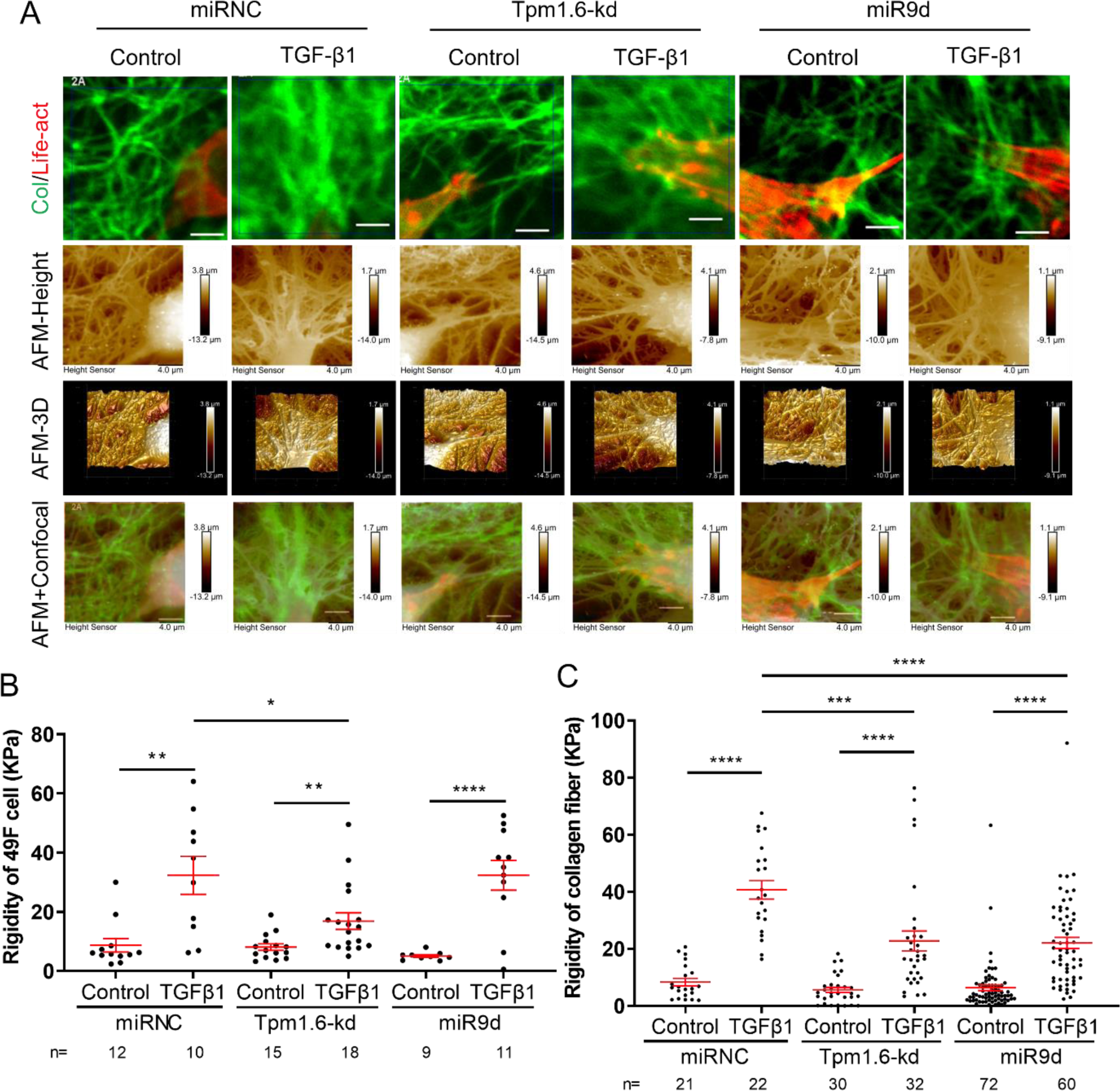
Silencing of Tpm1.6 reduces TGF-β1-induced augmentation of cell contractility and stiffness of collagen fibers. A-C. Cell contractility is measured by confocal plus atomic force microscope (AFM) system. LifeAct-expressing NRK-49F cells harboring biogenic RNA interference against Tpm1 were cultured on FITC-conjugated CG for 4 days with or without TGF-β1. Cells (LifeAct, red) and collagen fibrils (FITC, green) were visualized under confocal microscope and then subjected to co-axis atomic force microscopic analysis to assess the structure and rigidity of cells and peri-cellular collagen fibrils. AFM-Height images show the relative height information (μm) in topography. AFM-3D images display the surface morphology in three-dimensional presentation. AFM+Confocal indicates the merge of images between confocal microscope and AFM-Height. Scale bar = 20 μm. Quantification of rigidity of cells and peri-cellular collagen fibrils was shown in figure B and C, respectively. Statistical analysis was performed by two-way ANOVA.

### Tpm1.6-depletion triggers TGF-β1-induced collagen deformation via matrix metalloprotease (MMP)

To assess collagen degradation, we employed Cy3-conjugated collagen-hybridizing probe (CHP) which recognizes deformed collagen fibrils caused by traction force or enzymatic digestion (Bennink *et al*, 2018; Zitnay *et al*, 2017). Silenced control and Tpm1.6-kd cells were cultured on FITC-conjugated CGs for 24 h and followed by fixation and CHP incubation. Non-silencing control cells exhibited minimal degree of collagen aggregation and little CHP staining. TGF-β1 enhanced the collagen aggregation and triggered augmentation of CHP intensity which overlapped with collagen fibril. Inhibition of actomyosin activity by BB completely blocked TGF-β1-induced collagen aggregation and CHP intensity, whereas pan-MMP inhibitor (GM6001) did not alter TGF-β1-induced collagen aggregation and CHP intensity (Figure 6A, 6C), indicating that TGF-β1-indcued augmentation of CHP intensity is predominantly contributed by cell contractility. Interestingly, Tpm1.6-kd did not alter TGF-β1-increased collagen aggregation and CHP intensity. BB markedly blocked TGF-β1-induced collagen aggregation and CHP intensity, whereas GM6001 treatment completely alleviated TGF-β1-induced CHP intensity, suggesting that TGF-β1-induced collagen deformation in Tpm1.6-kd cells is mediated by activation of MMPs (Figure 6B, 6C).

**Figure 6.**
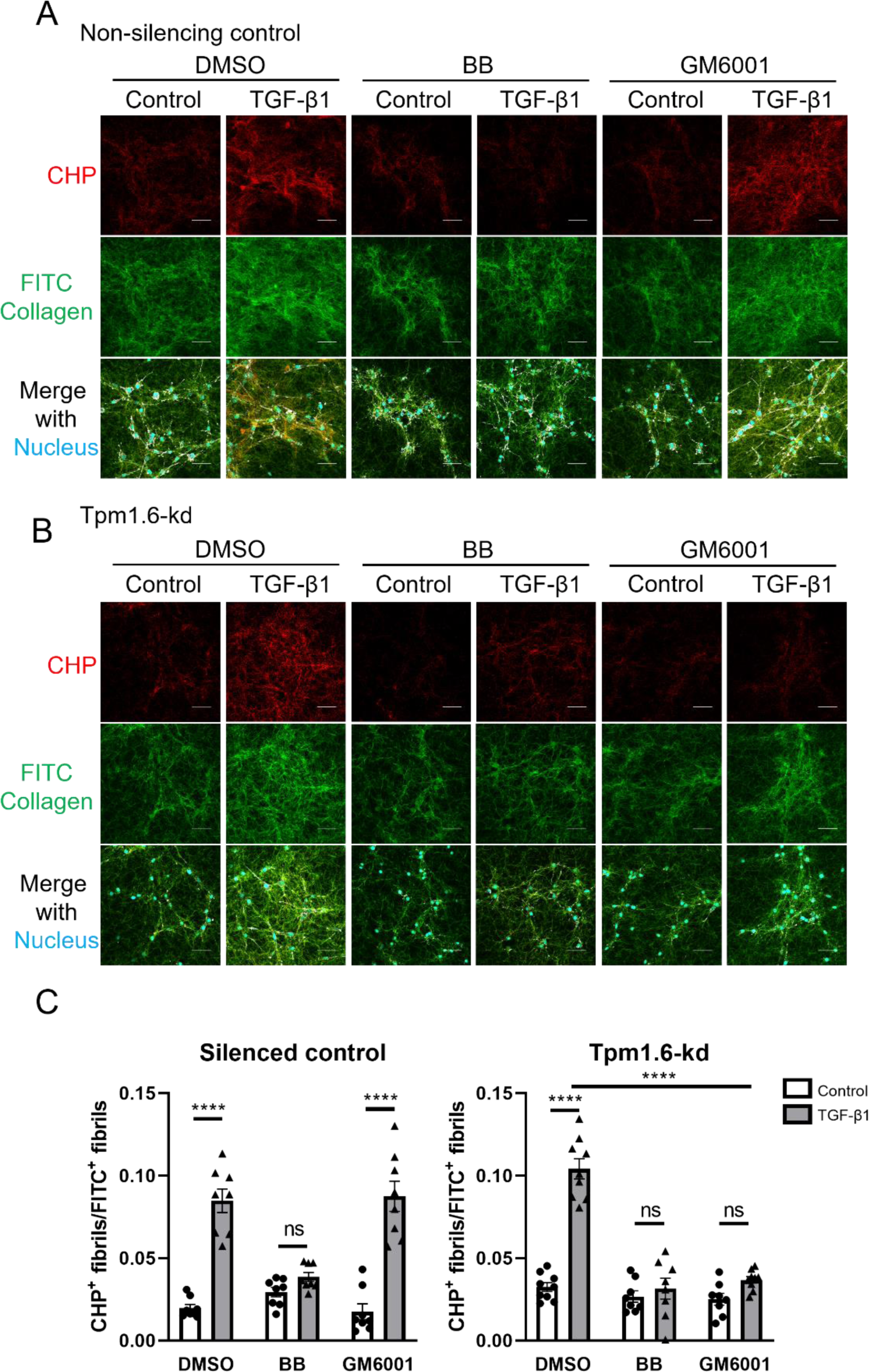
The collagen deformation caused by Tpm1.6-kd fibroblasts upon TGF-β1 stimulation is mediated by MMPs. Non-silenced and Tpm1.6-silenced NRK-49F cells were cultured on FITC-conjugated collagen gel with or without TGF-β1 for 24 h and co-treated with actomyosin blocker (BB-blebbistatin) or pan-MMP inhibitor (GM6001). The culture was fixed and collagen gels were incubated with Cy3-collagen hybridizing probes (CHP) for 2 h to detect the deformed collagen fibrils caused by cell traction force or enzymatic digestion. A-C. Fluorescent images show the deformed collagen fibrils (Cy3-CHP, red) and total collagen fibrils (FITC, green) in culture of non-silenced (A) or Tpm1.6-silenced (B) NRK-49F cells, and the quantification of intensity of CHP-positive fibrils vs FITC-collagen fibrils was shown in figure C. TGF-β1-induced deformation of collagen fibrils by Tpm1.6-silenced NRK-49F cells is completely inhibited by GM6001, indicating that MMP-triggered collagen degradation is involved. n=8. Scale bar = 20 μm. Statistical analysis was performed by two-way ANOVA.

### Tpm1.6-depletion triggers TGF-β1-induced collagen degradation via MMP9-enriched α-SMA aggregates

To evaluate whether podosome formation is involved in active collagen degradation capability in Tpm1.6-kd fibroblasts. Silenced control and Tpm1.6-kd fibroblasts were cultured on CGs for 24 h with or without TGF-β1, and the canonical podosome structure was subsequently analyzed by cortactin- and pholloidin-staining. Control cells displayed little amount of podosome structure. TGF-β1 or Tpm1.6-depletion slightly increased the podosome formation, but markedly enhanced podosome dots and podosome belt in TGF-β1-treated Tpm1.6-kd cells. (Figure 7A-upper). When the cultured cells were co-stained with α-SMA and cortactin, we have very surprising observations. In control cells, TGF-β1 increased α-SMA level, but the expression of α-SMA was exclusive of cortactin staining. In Tpm1.6-kd cells, we observed the presence of α-SMA dots. Interestingly, TGF-β1 triggered a marked increase of α-SMA dots with a larger size near the cell membrane (Figure 7A-lower, 7B). The presence of α-SMA dots were verified by another targeted sequence of exon 2b of Tpm1.6 (Appendix Fig S4C). These α-SMA dots showed few co-staining with cortactin, suggesting that they are not canonical podosome (Figure 7A).

**Figure 7.**
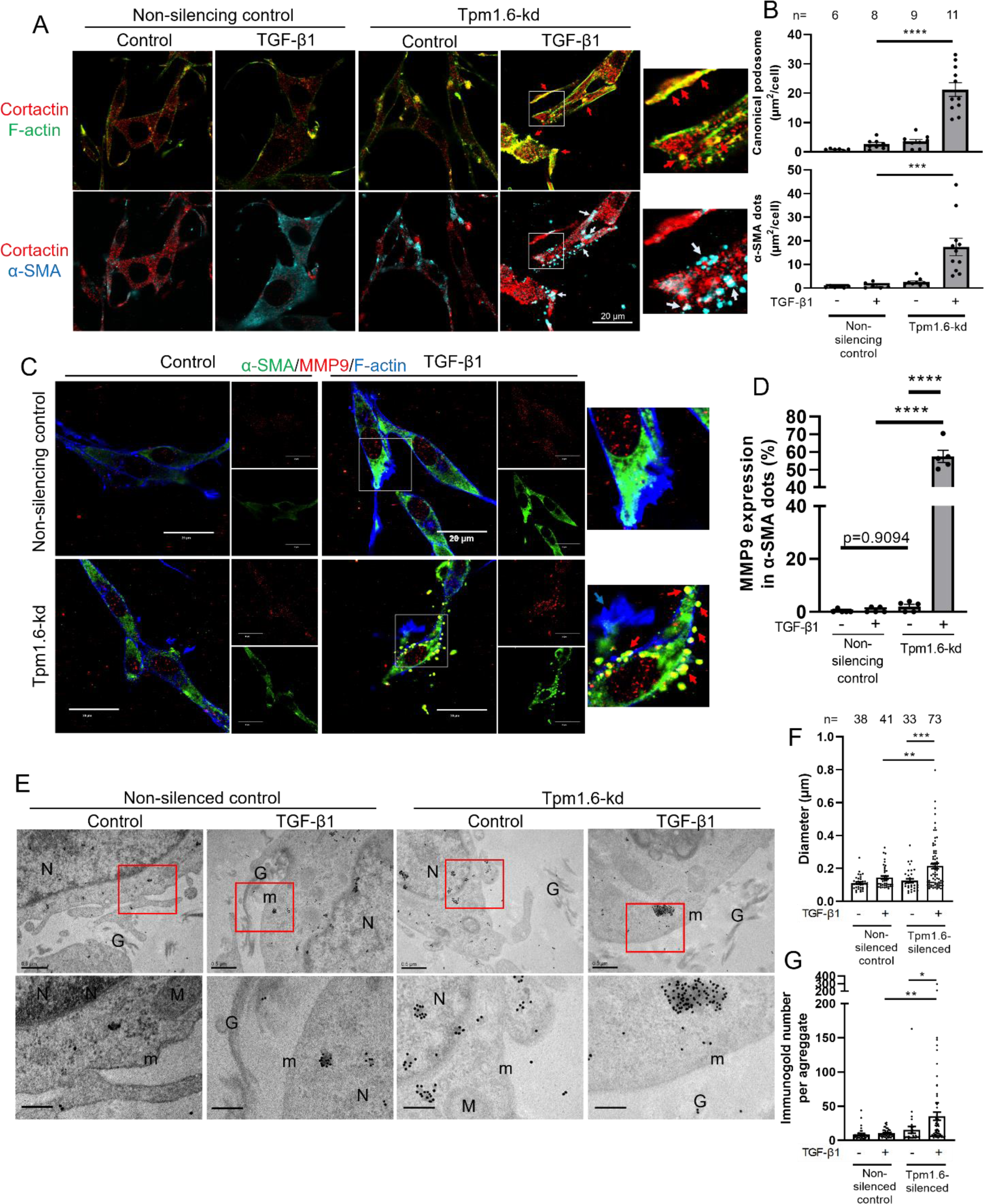
Silencing of Tpm1.6 by exon 2b-targeted RNA interference triggers the formation of canonical and non-canonical podosome upon TGF-β1 induction. Non-silenced and Tpm1.6-silenced NRK-49F cells were cultured on CG with or without TGF-β1 for 24 h and, the cultured cells underwent immunofluorescence studies and transmission electron microscope (TEM) inspection. A. Canonical podosomes were represented by cortactin (red), phalloidin (green) staining; non-canonical podosomes were shown by α-SMA dot (cyan). Tpm1.6-silenced NRK-49F cells displayed more canonical and non-canonical podosomes with TGF-β1 induction. Scale bar = 20 μm. B. Quantitative results show the size (μm^2^) of canonical podosomes and α-SMA dots in non-silenced and Tpm1.6-silenced NRK-49F cells. Statistical analysis was performed by two-way ANOVA. C. Fluorescent images of the localization of α-SMA (green) and MMP9 (red) show that there is MMP9 enrichment in non-canonical podosomes in Tpm1.6-silenced NRK-49F cells. Scale bar = 20 μm. D. Quantitative results of the percentage of MMP9-positive area in α-SMA dots in non-silenced and Tpm1.6-silenced NRK-49F cells with or without TGF-β1. n=5. Statistical analysis was performed by two-way ANOVA. E. The cell-collagen scaffolds were embedded by Spurr resin after fixation and dehydration. The embedded scaffolds were sectioned with a thickness of 70 nm and then incubated with α-SMA antibody and immunogold-conjugated secondary antibodies. The images of TEM showed the detailed structure of cell-matrix adhesion site and α-SMA-positive aggregates. G – CG; N – nucleus; m – cell membrane; M – mitochondria. Scale bar = 0.5 μm (upper) and 0.2 μm (lower). F, G. Quantitative results show the α-SMA-enriched aggregate size and the number of captured gold per aggregate. Statistical analysis was performed by two-way ANOVA.

To explore the possible role of α-SMA dots, we co-stained MMP9. MMP9 was expressed at a low level in control cells. TGF-β1 increased the levels of MMP9, which was not co-stained with α-SMA. Tpm1.6-kd did not alter the expression of MMP9, whereas TGF-β1 not only increased the MMP9 levels but also markedly induced colocalization of MMP9 and α-SMA dots (Figure 7C, 7D). These findings indicate that Tpm1.6 depletion facilitates the formation of MMP-containing α-SMA dots upon TGF-β1. To visualize the detailed structure of α-SMA dots, we employed transmission electron microscope (TEM) and α-SMA-targeted immunogold. Control and Tpm1.6-kd cells were cultured on CGs for 24 h and followed by fixation, embedding, sectioning, and subjected for TEM examination. The diameter of α-SMA aggregates at the cell-matrix adhesion site in control cells was around 0.1 μm, and TGF-β1 did not alter the size. In Tpm1.6-kd cells, the size of α-SMA aggregates was not altered, but TGF-β1 induced larger α-SMA aggregates (Figure 7E-G), consistent with immunofluorescence studies. Because such α-SMA aggregates are enriched in MMP9, but devoid of cortactin, we further test whether they encompass capability of collagen degradation.

To assess the collagen-degraded capability in cells, we analyzed the levels of degraded collagen protein released into culture media. Control and Tpm1.6-kd fibroblasts were cultured on CGs for 24 h, and then CGs were either released or treated with GM6001 for 48 h. The protein components in culture media were precipitated and analyzed by Western blotting. Only collagen monomer and polymer were present in culture media derived from control cells. TGF-β1 induced an increase in levels of collagen polymer (above 100 kDa). Tpm1.6-kd did not affect collagen release in culture media; however, TGF-β1 markedly increased the levels of collagen degradation products released into the media (Figure 8A, 8F). To deplete physical force of CG substrates, we released the attachment between gel and culture dish. The release of CG for two days did not altered collagen levels, but reduced TGF-β1-induced collagen polymer in culture media from control cells. Tpm1.6-depletion did not alter the release of CG-induced collagen levels, but markedly trigger TGF-β1-induced collagen degradation as reflected by reduced level of monomer and augmentation of the smallest collagen fragment in culture media (Figure 8A, 8F). The Western blotting results were verified by another type I collagen antibody (Appendix Fig S5A). GM6001 did not alter the pattern of collagen degradation in the culture media in control and Tpm1.6-kd cells, but completely blocked TGF-β1-induced collagen degradation in Tpm1.6-kd cells (Figure 8B, 8G). To examine the function of MMP9-containing α-SMA aggregates in collagen degradation, we employed pharmacological MMP9 inhibitor and biogenic silencing of α-SMA (shSMA). Neither inhibition of MMP9 nor depletion of α-SMA altered the collagen degradation pattern in culture media in control and Tpm1.6-kd cells. However, either inhibition of MMP9 or depletion of α-SMA completely blocked TGF-β1-induced collagen degradation only in Tpm1.6-kd cells (Figure 8C, 8D, 8H, 8I). These results indicated that α-SMA aggregates serve collagen degradation in Tpm1.6-kd fibroblasts upon TGF-β1 induction. Additionally, depletion of typical podosomal protein (siCortactin) also attenuated TGF-β1-induced collagen degradation in Tpm1.6-kd cells (Figures 8E, 8J). Based on immunofluorescence studies and Western blotting results, TGF-β1 predominately triggers collagen degradation in Tpm1.6-kd cells via MMP9-containing α-SMA aggregates. These findings reveal that a new structure contributes to the collagen degradation.

**Figure 8.**
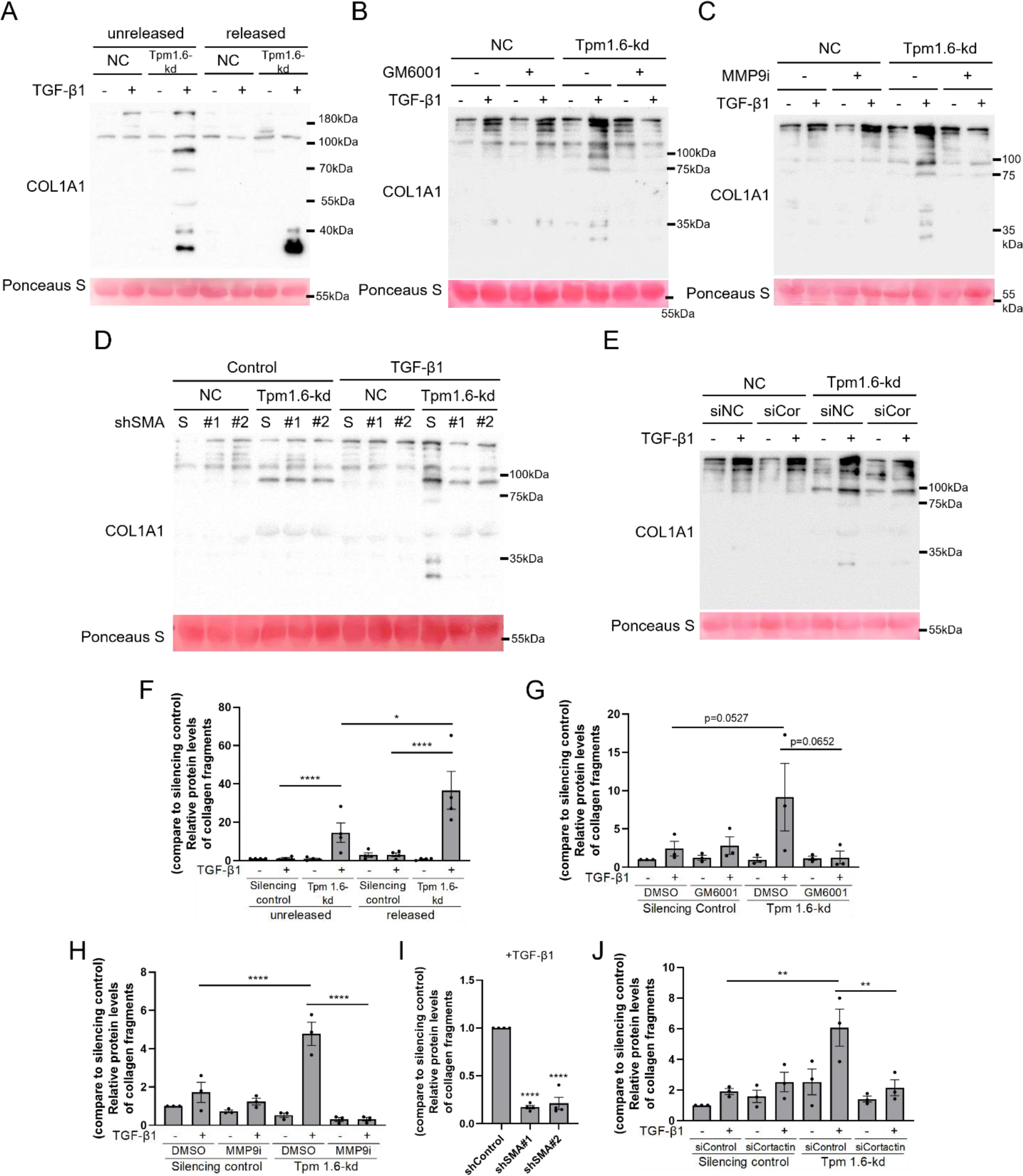
Suppression of Tpm1.6 promotes collagen digestion upon TGF-β1 stimulation via induction of non-canonical podosome. Non-silenced and Tpm1.6-silenced NRK-49F cells were cultured on CG for 24 h and then the CG was either released (A) or were treated with MMP inhibitors (GM6001(B) or MMP9i (C)) for 48 h with or without TGF-β1. The culture media was harvested for Western blotting analysis to detect collagen degradation. A, F. Western blotting results of the culture media collected from cells cultured on unreleased or released CG. Tpm1.6-silencing markedly increased collagen fragmentation (smaller than the size of monomer) upon TGF-β1 induction. The release of CG to reduce traction force triggers the collagen degradation effect, and the quantification is shown in (F). n=3. Statistical analysis was performed by two-way ANOVA. B, C, G, H. Western blotting results to show the effects of GM6001 (B) or MMP9 inhibitor (C) on TGF-β1-induced collagen degradation in Tpm1.6-silenced cells cultured on unreleased CG. Both GM6001 and MMP9 inhibitor completely suppressed TGF-β1-induced CG degradation in Tpm1.6-silenced cells, and the quantification is shown in (G) and (H), respectively. n=3. Statistical analysis was performed by two-way ANOVA. D, E, I, J. Non-silenced and Tpm1.6-silenced NRK-49F cells were transfected with shSMA (D) or siCortactin (E) and cultured on unreleased CG with or without TGF-β1 for 48 h. The culture media were collected for Western blot analysis. Inhibition of α-SMA (non-canonical podosome) or cortactin (canonical podosome marker) markedly reduced TGF-β1-induced collagen degradation exerted by Tpm1.6-silenced cells, and the quantification is shown in figure I and J, respectively. n=3. Statistical analysis was performed by one-way ANOVA (I).

## Discussion

In our study, we investigated myofibroblast activation under an *in vivo*-like environment to mimic the initial stage of fibrosis progression. Only fibroblasts were activated into myofibroblasts, which facilitated by actomyosin-associated CUs. We found that actomyosin contractility plays a crucial role in myofibroblast activation, suggesting a reciprocal relationship between the remodeled matrix and fibroblast activation. Kramann’s study also supported the notion that fibroblasts are the primary precursors of myofibroblasts in renal fibrosis (Kuppe *et al*., 2021). Thus, fibroblasts likely serve as the initial precursor population for myofibroblast activation through actomyosin-mediated matrix remodeling during the onset of fibrosis progression.

Targeting exon 2b and 9d for Tpm1.6 depletion displayed different outcomes in myofibroblast activation and collagen degradation. Tpm1.6 is the isoform associated with stress fibers, suggesting distinct role among isoforms in the phenotypic switch (Gateva *et al*., 2017). Figure 3G revealed that both exon 2b and exon 2a were silenced in miR9d cells, suggesting alterations in other isoforms (Tpm1.3/1.4) which may cause minor impact on phenotypic change in miR9d cells. The presence of shared exons within isoforms poses a limitation in recognizing Tpm isoforms and targeted therapy. Nevertheless, we employed Sanger sequencing and Taqman probes to identify the specific Tpm. Our findings highlight Tpm1.6 as the major isoform in renal fibroblasts with TGF-β1-specific regulation, underscoring the significance of investigating the function of Tpm1.6. Overall, Tpm1.6 acts as the major fibroblast isoform controlling the phenotypic switch in renal fibroblasts.

As shown in Figure 1, cell contractility and podosome-formation are mutually exclusive. Tpm1.6-depletion blocks TGF-β1-induced cell contractility, while simultaneously promoting collagen degradation through α-SMA aggregates in response to TGF-β1. Additionally, reduction of external force by gel release enhances TGF-β1-induced collagen degradation in Tpm1.6-silenced cells (Figure 8A). Such findings highlights that the phenotypic change is controlled by Tpm1.6 and mechanical properties of substratum.

The function of Tpm is to stabilize actin filament. It is likely that specific isoform of Tpm may interact with diverse actin isoform. Tpm1.6-depletion significantly changes TGF-β1-induced α-SMA expression pattern via reduction of stress fiber-associated α-SMA and induction of membrane-associated α-SMA aggregates. Tpm1.6 seems to display more affinity with contractile α-SMA. Actin depolymerizing factor (ADF) and filaminm have been reported to compete with Tpm for the association of actin filaments with isoform-specificity (Bryce *et al*, 2003; Kovar *et al*, 2011; Ono & Ono, 2002). Tpm regulates the activity of myosin motors and the intracellular sorting of myosin II depending on isoforms (Coulton *et al*, 2010; Fanning *et al*, 1994; Strand *et al*, 2001; Tojkander *et al*, 2011). Tropomodulin maintains actin homeostasis through distinct affinities for Tpm isoforms (Kostyukova & Hitchcock-DeGregori, 2004; Kumari *et al*, 2020; Uversky *et al*, 2011). Taken together, distinct Tpm/actin affinity and other regulatory proteins may be involved in phenotypic change from contractile to matrix-degrading machinery exerted by Tpm1.6-depletion.

TGF-β superfamily encompasses a diverse group of signaling molecules, including TGF-β1 and BMP, which activate distinct smad signaling pathways. TGF-β1 predominantly triggers p-smad2/3, while BMP promotes p-smad1/5/8 pathway. Recent studies revealed the activation of non-canonical smad1/5/8 axis upon TGF-β1, suggesting an alternative signaling mechanism (Jann *et al*, 2020; Ota *et al*, 2013). Additionally, TGF-β superfamily was implicated in osteoclast maturation, which is known for high collagen-degraded capability. This degraded phenotype bears resemblance to the findings observed in Tpm1.6-depleted fibroblasts. The involvement of smad1/5/8 signaling in osteoclast maturation may contribute to collagen degradation; therefore, TGF-β1-smad1/5/8 axis may be responsible for degraded phenotype in Tpm1.6-depleted fibroblasts. To fully comprehend the relationship between different smad regulations and collagen degradation, further investigation is warranted.

Our findings demonstrated that Tpm1.6 is a specific isoform in renal fibroblasts which play pivotal roles in determination of contractile phenotype or collagen degradation upon TGF-β1. Tpm1.6 contributed to cell contractility, and the depletion switched contractile phenotype to matrix-degrading phenotype upon TGF-β1. Because the α-SMA aggregates are highly colocalized with MMP9, but not cortactin, such collagen digesting structure may be a kind of non-canonical podosome. The matrix-degrading phenotype is mediated by non-canonical podosomes. Such findings point out the crucial role of Tpm1.6 in initial stage of renal fibrosis. These findings demonstrated a novel way for prevention of fibrosis progression and potential fibrosis therapy.

## Author Contributions

CLW and MJT designed research; CLW, GHL, BYC and TTC performed research; YKW contributed analytic tools; CLW analyzed data; CLW, and MJT wrote the manuscript in consultation with CHK, and TYW.

## Acknowledgements

The authors appreciate for members in National Cheng Kung University Medical College Core Research Laboratory for technical experimental help. Funding was provided by the Ministry of Education (Higher Education SPROUT Project: International Center for Wound Repair and Regeneration), Ministry of Science and Technology (MOST 109-2634-F-006-021, MOST 110-2634-F-006-018, MOST 111-2634-F-006-009 to MJT; MOST 109-2320-B-006-019 to CHK).

## Author contributions

Ching Lung Wu and Ming Jer Tang designed research; Ching Lung Wu, Gang Hui Lee, Bo Yu Chen and Ting Ting Chan performed research; Yang Kao Wang contributed analytic tools; Ching Lung Wu analyzed data; Ching Lung Wu and Ming Jer Tang wrote the manuscript in consultation with Cheng Hsiang Kuo, and Tzyy Yue Wong.

